# Bayesian Estimation of Co-occurrence *Affinity* with Dyadic Regression

**DOI:** 10.1101/2024.01.16.575941

**Authors:** Arthur Newbury

**Affiliations:** Centre for Ecology & Conservation, College of Life and Environmental Sciences, University of Exeter, Penryn, Cornwall, TR10 9FE, UK

**Keywords:** Bayesian, biogeography, co-occurrence, functional traits, phylogeny

## Abstract

1. Estimating underlying co-occurrence relationships between pairs of species has long been a challenging task in ecology as the extent to which species co-occur is partially dependent on their prevalence. While recent work has taken large steps towards solving this problem, the next question is how to assess the factors that influence co-occurrence.
2. Here, I show that a recently proposed co-occurrence metric can be improved upon by assigning Bayesian priors to the latent co-occurrence relationships being estimated. In the context of analysing the factors that affect co-occurrence relationships, I demonstrate the need for a generalised linear model (GLM) that takes raw data (co-occurrences and species prevalence) not derived quantities (co-occurrence metrics) as its data. Next, I show the form that such a GLM should take in order to perform Bayesian inference while accounting for non-independence of dyadic matrix data (e.g. distance and co-occurrence matrices).
3. I then present 3 example analyses to highlight the types of scientific questions these methods can help answer, using existing data sets - measuring the effects of trait dissimilarity among dung beetle species and relatedness between ant species on co-occurrence, and constructing co-occurrence networks of bacteria found in cystic fibrosis patient sputum samples.
4. Finally, I present the software package CooccurrenceRegression.jl, which provides a straightforward interface for researchers to put these methods into practice.

## 1 Introduction

The analysis of patterns of co-occurrence between taxa is an important and active area of ecological research (Gotelli and McCabe 2002; Barberán et al. 2012; Williams, Howe, and Hofmockel 2014; Kraan, Thrush, and Dormann 2020; Zhu et al. 2023). Mainali et al. (2022) have recently shown that of the numerous ways of measuring co-occurrence relationships between species pairs (reviewed in (Mainali et al. 2022)) the correct and unbiased method is to make use of Fisher’s noncentral hypergeometric distribution, as suggested by Veech (2013) and Griffith et al. (2016). Mainali et al. (2022) derived a co-occurrence metric called *affinity* (or *α*) based on this distribution. This is a significant step forward in the analysis of cooccurrence relationships. There are still several challenges remaining, however. For instance, when species pairs of very low or very high prevalence are analysed with this method, they will often be assigned very high or low affinity scores, but with low confidence (high p-value and wide confidence interval). For downstream analysis then the researcher may (i) treat all data points as equal, (ii) remove data above some high p-value threshold or (iii) devise a scheme to weight data appropriately. (i) wastes information and can yield misleading results. (ii) again, wastes information and could severely bias results depending on the reasons for the differences in prevalence/p-values. If an appropriate, unbiased scheme for (iii) should be devised, then this would be a welcome development. However, adopting a Bayesian approach that builds upon the work of Mainali et al. (2022) not only yields more accurate estimates of the *affinity* between species, but also naturally propagates uncertainty through the analysis, i.e., it accounts for the differing levels of confidence we have about the co-occurrence relationships between different species pairs. In the present work I illustrate this point and provide the model description and code to use this Bayesian method in practice.

A significant advantage of this framework is that it allows co-occurrence data to be analysed as response data in a Bayesian general linear model (GLM). That is, by supplying species occurrence data as the prevalence of individual species, the number of times a given species pair co-occur and the total number of sites considered, it is now no more complicated to construct a regression model (with a Fisher’s noncentral hypergeometric likelihood function) than it would be to perform binomial regression with count data. Similarly, this method makes the best use of all available information and weighs data points appropriately. Thus, just as it is not appropriate to convert count data to proportions and conduct linear regression, it is no longer best practice to summarise co-occurrence data as point estimates for linear regression.

## 2 Materials and Methods

### 2.1 Simulation data

In the following, maximum likelihood estimates of co-occurrence affinity were obtained using the R (R Core Team 2014) package `co-occurrenceAffinity` (Mainali et al. 2022). All other analyses and visualisations were carried out in Julia (Bezanson et al. 2017). All models were constructed in the probabilistic programming language `Turing` (Ge, Xu, and Ghahramani 2018) with MLE estimates of regression coefficients fit to data using the Nelder-Mead method (Nelder and Mead 1965) and MAP estimates of *α* fit using L-BFGS (Liu and Nocedal 1989) implemented in `Optim` (Mogensen and Riseth 2018) and results visualised in `Makie` (Danisch and Krumbiegel 2021).

#### Maximum *a posteriori* vs maximum likelihood pairwise affinity estimates

Mainali et al. (2022) show that the log of the odds ratio term in Fisher’s noncentral hypergeometric distribution (a quantity they term *α*) can be used to appropriately describe the extent to which two species will tend to co-occur more or less than would be expected based just on the prevalence of the species. They go on to propose that the maximum likelihood estimate of this parameter 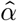 or *affinity* should be used as a pairwise co-occurrence metric. However, Maximum likelihood estimates can yield values of positive or negative infinity. This causes difficulties for downstream analyses (one cannot do something as simple as calculating the mean of a set containing infinite values). Furthermore, we know *a priori* that an infinitely large or small affinity is not sensible for most cases of interest to ecologists. Bayesian analysis uses prior knowledge to avoid the estimation of physically or biologically implausible values. Mainali et al. (2022) reassign these infinite estimates an absolute value of log(2*N* ^2^), where *N* is the total number of sample sites (from an argument made based on the Jeffreys’ prior for the beta distribution). While this figure comes from sound argument, there are at least two problems with this approach. Firstly, not all data are treated the same way, i.e., no regularisation is applied to finite affinity estimates, only to these extreme values. Secondly, the value log(2*N* ^2^) is only a function of *N*, not the species prevalence. Thus, it is not influenced by our actual state of knowledge about the species in question.

To make these ideas more concrete and show the practical implications, I simulated species pairs using the affinity model. For each pair there are *N* = 30 sites they can inhabit, species *A* has a prevalence *mA* and species *B* has prevalence *mB*. The number of sites at which they co-occur *k* was drawn from a Fisher’s noncentral hypergeometric distribution

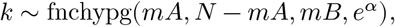

with 10 draws per combination of *mA* and *mB* for each of 41 different values of *α*. Then, given the values for *N, k, mA* and *mB* I estimated *α* using two methods. Firstly, I used the original maximum likelihood estimate (MLE) of Mainali et al. (2022). Next, I obtained maximum *a posteriori* (MAP) estimates with a Gaussian prior N(0, 3) for *α*. Note that these are not strongly regularising priors as when exponentiated in the likelihood function a standard deviation of 3 ≈ 20 and two standard deviations 6 ≈ 403 which is a very large odds ratio for most applications. In order to compare the two methods, for each combination of *mA* and *mB* I calculated the deviation between the inferred and ground truth *α* values as the root mean squared error (RMSE).

#### Regression analysis

Often, we do not simply wish to report co-occurrence relationships, but to measure how they change with some other variables of interest. For many types of data there are well understood and regularly used probability distributions which can be used in Bayesian and frequentist GLMs. For co-occurrence data this has not been the case. Given the issues with deriving point estimates highlighted above it is unlikely that simply fitting a linear model to such point estimates of co-occurrence affinity will yield reliable results. Thus, rather than supplying a second inference model with uninformative point estimates from a previous model, we can provide our regression model with all the data on species prevalence and co-occurrences instead.

To demonstrate the impact of this I simulated 41 sets of predictor data 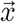, each consisting of 30 draws from N(0, 1). Affinity values were generated by multiplying the predictor data by a regression coefficient *β*. For each affinity value - species prevalence *mA* and *mB* values were chosen randomly between 1 and 29 and a *k* value was drawn from the Fisher’s noncentral hypergeometric distribution as above. For each generated data set pairwise affinity values were estimated by the MAP and MLE methods. Then linear regression analysis was conducted on these point estimates 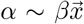. Additionally, I obtained a maximum likelihood estimate of a GLM of the form

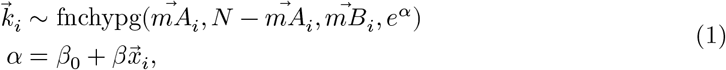

where 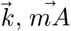 and 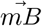 are vectors containing the values of *k, mA* and *mB* respectively and *β*_0_ is the intercept.

#### Dyadic regression

In many if not most cases when working with co-occurrence data it will be in the form of a square co-occurrence matrix similar to the distance and dissimilarity matrices used to record e.g. phylogenetic distances between species or community dissimilarities between sample sites. As with these other types of matrices, if we wish to perform regression analysis treating each entry in the matrix as data point, we must account for non-independence of data coming from the same row or column, e.g. same site, species etc.. To deal with this we can include a random effect *λ* for each species (Clarke, Rothery, and Raybould 2002; Gompert et al. 2014). Thus, whereas the GLM above contains the term

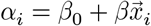

we would now have

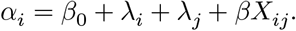

Where *X* is a (possibly dissimilarity or distance) matrix in which element *X*_*ij*_ is a quantity of interest relating species *i* to species *j*. Given a similarly arranged matrix *K* which holds the *k* values for all species pairs and a vector 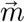 containing species prevalence, the dyadic GLM becomes

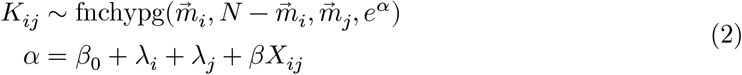

for each index *ij* in either the lower or upper triangle of *K*. While Maximum likelihood estimates of the *λ, β* and *β*_0_ parameters can be obtained, we still need a way to properly account for our uncertainty in our estimates. Bayesian inference provides an intuitive framework for this, grounded in probability theory. Thus, we can construct a Bayesian GLM by assigning priors to the unknown parameters. Assuming Gaussian priors for all parameters gives

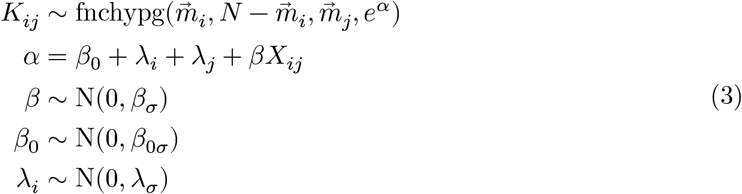

where *β*_*σ*_, *β*_0*σ*_ and *λ*_*σ*_ are to be chosen according to the specifics of the system being analysed. This forms the basic model structure used for analyses in the examples with real data.

### 2.2 Examples with real data

In the following examples all explanatory variables were standardised to have mean 0 and standard deviation 1, in order to make results comparable. Inference was performed via Markov chain Monte Carlo (MCMC), using the No U-turns (NUTS) sampler (Homan and Gelman 2014) implemented the Turing probabilistic programming language (Ge, Xu, and Ghahramani 2018), with 8 chains of 1000 iterations in each case. Point estimates are reported with 95% credible intervals (CI) calculated from quantiles of MCMC samples and probability of direction (PD), the posterior probability that the reported effect is in the estimated direction.

#### Change in co-occurrence affinity owing to trait dissimilarity amongst dung beetles

For the first example I used previously published occurrence data for dung beetles along an altitudinal gradient at Serra do Cipó, State of Minas Gerais, Brazil (Nunes et al. 2016). The data set includes occurrence data for 56 beetle species, across 7 sites, ranging in elevation from 800 to 1400 meters above sea level, as well as mean biomass (dried weight) in grammes and the functional guild for each species. In order to assess how functional trait dissimilarity affected co-occurrence affinity we employed 3 different regression models 1) combining the two functional traits into a single Gower’s distance (Gower 1971); 2) including both traits and their interaction as explanatory variables; 3) focussing on the effect of biomass *within* functional guild. Thus, model 1 was identical in form to equation Equation 3

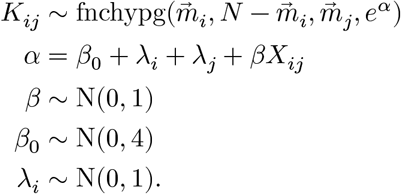

Model 2 was

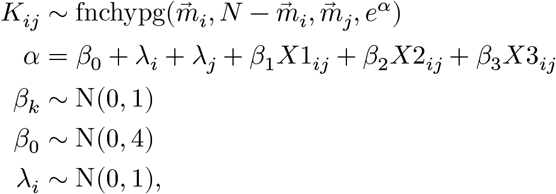

where *X*3 = the element-wise product *X*1 ⊙ *X*2. Model 3 was

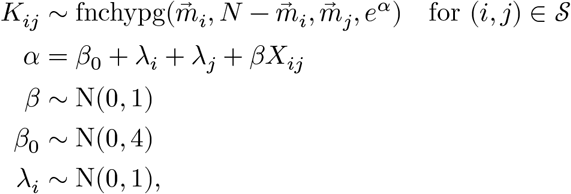

where 𝒮 = {(*i, j*) : *g*_*i*_ = *g*_*j*_} is the set of indices denoting species pairs from the same functional guild.

#### Change in co-occurrence affinity owing to trait dissimilarity amongst ants

For the second example I used two data sets, one containing global ant species occurrence data and phylogenetic tree (Economo et al. 2018) and containing ant occurrence data and trait measurements from two nature reserves near Hong Kong (Wong et al. 2021). The trait data consisted of species mean values for body size, leg length, head width, mandible length, pronotum width and scape length. In order to analyse the relationship between trait dissimilarity and co-occurrence affinity I first subset the data to work with only those species that occurred in both data sets, so that I had trait measurements and occurrence data at both scales for all of them. Separate models were then used for the two geographic scales, following the same basic model structure as above for the global data set

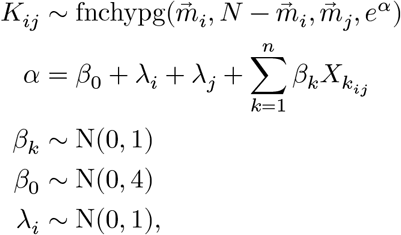

where here *n* = 6 traits. However, the the local scale data set used dat from two separate nature reserves and from two types of sites: those where invasive species *Solenopsis invicta* was present and those where it was absent. I pooled the data from these four site types, with random intercepts for each, giving the model

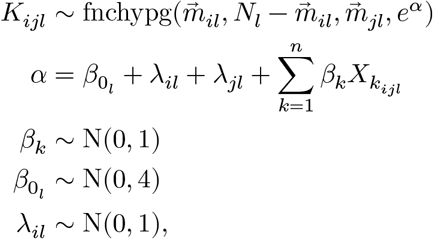

In order to determine the relationship between phylogenetic distance and co-occurrence affinity I used 3 models: 1) measuring the relationship at genus level; 2) at species level with a single regression coefficient; 3) at species level with a hierarchical model. The phylogenetic tree was read and traversed using phylo.jl (Reeve, Borregaard, and Harris 2024). Genus level phylogenetic distances were calculated as the tree distance between most recent common ancestor node (MRCA) of one species and the MRCA of another. Prior to conducting the species level analysis, I removed all genera with less than 25% unique phylogenetic distances. This was to reduce any bias introduced by species having artificially low phylogenetic distance due to lack of resolution in the tree. 61 genera remained for analysis. These analyses were conducted only across the global geographic scale, as there was insufficient phylogenetic data to calculate distances between many of the species at the local scale.

Analysing the effect of phylogenetic distance on co-occurrence at the genus level took an identical form to Equation 3 with same priors as model 1 of the dung beetle analysis. Similarly, the pooled species level model was identical to model 3 of the dung beetle analysis. The hierarchical model was given by

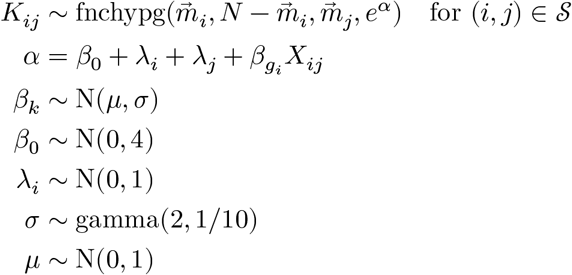

where 𝒮 = {(*i, j*) : *g*_*i*_ = *g*_*j*_} is the set of indices denoting species pairs from the same genus and *g* is a vector of indices mapping each species to it’s genus.

#### Bacterial co-occurrence networks derived from cystic fibrosis patient sputum samples

To demonstrate another use for Bayesian estimation of co-occurrence affinity I used microbiome data from cystic fibrosis patient sputum samples (Quinn et al. 2019) to construct co-occurrence networks. The data originally consisted of 4148 unique sequences of 100 nucleotides of the v4 region of the bacterial 16S rRNA (primers: 515f GTGCCAGCMGCCGCGGTAA; 806r GGACTACHVGGGTWTCTAAT). All sequences were identified to genus level using the RDP classifier (Wang et al. 2007) implemented in AssignTaxonomy.jl. Prior to analysis, I removed all sputum samples which were outliers in terms of read depth (8000 < read depth < 16000) to reduce the impact of differences in read depth on affinity estimates e.g. rare taxa co-occurring in bigger samples. After combining occurrence data relating to sequences from the same genera and limiting the data to a single sample from any one patient, the final data set consisted of a presence/absence matrix of 287 genera across 62 patients.

The affinity of each genus pair was inferred separately:

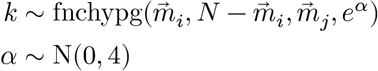

The same computational methods were used as for all the other examples, with the exception that I used only a single MCMC chain for each genus pair. Results were visualised both as an adjacency matrix an as networks.

## 3 Results

### 3.1 Simulation data

#### Maximum *a posteriori* vs maximum likelihood pairwise affinity estimates

Fig 1 shows that only when both *mA* and *mB* were equal to 15 was the RMSE approximately equivalent between MLE and MAP methods. Whenever one or both species had a high or low prevalence, and particularly as the absolute value of *α* became larger, the MLE method produced very poor estimates, and the extreme estimates were always the same log(2*N* ^2^) = 7.496. By contrast, for the MAP values, the prior provides regularisation which can be overcome by increasing confidence in the data, which is a function of *mA* and *mB*. Thus, the models *best guess* when *mA* = 15, *mB* = 5 and *k* = 5 is higher than the equivalent situation when *mB* = 1 and *k* = 1. Neither of these methods is perfect, however. When asked for one, a model will give you it’s best guess point estimate, but we can make better use of the data we have collected if we can utilise not only the point estimates but also our uncertainty around them.

**Figure 1:**
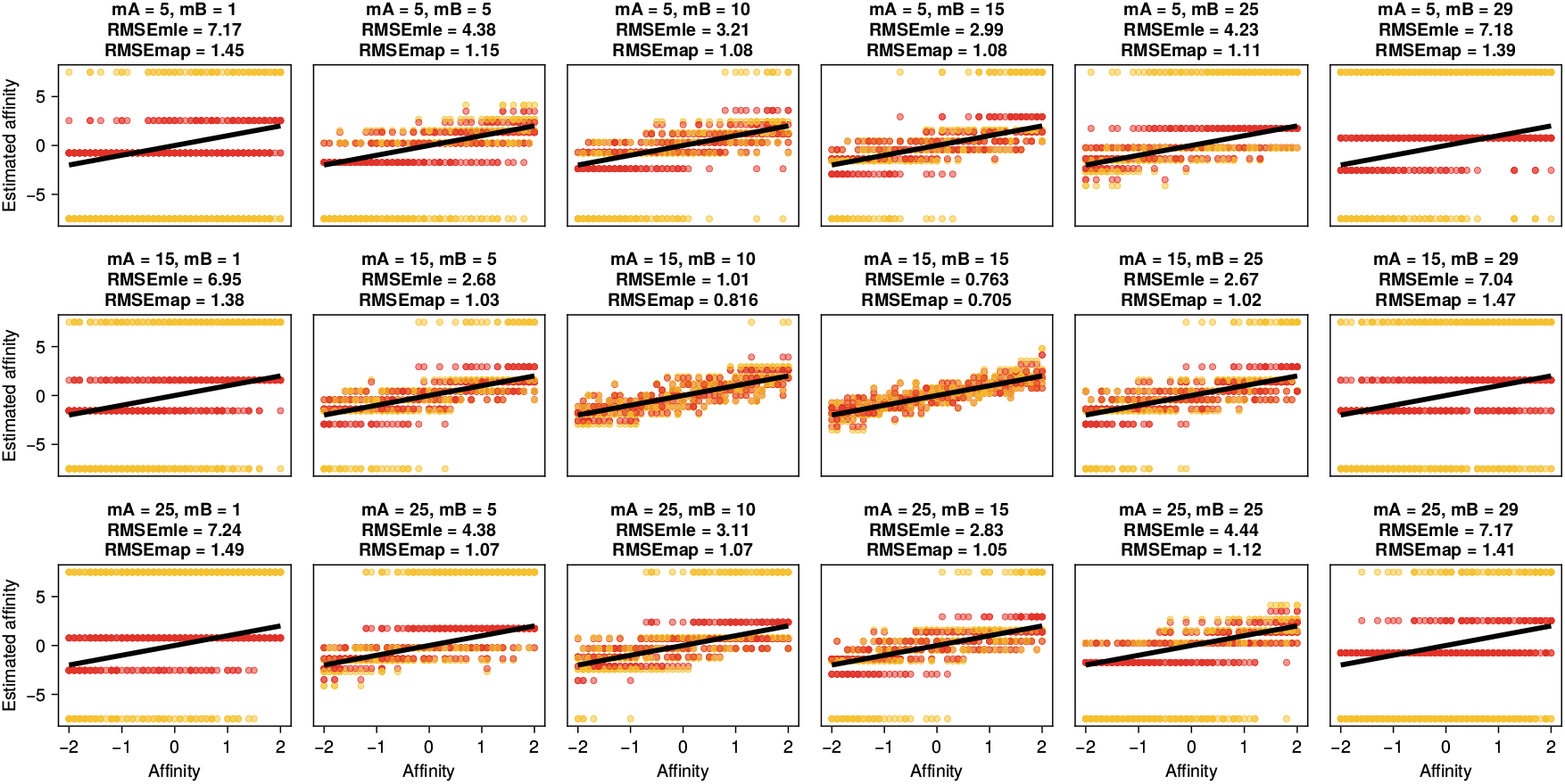
Actual and estimated affinity values for a range of species prevalences. Yellow points are estimates using the original maximum likelihood method and red points are maximum *a posteriori* estimates. Black lines indicate the actual affinity values used to generate the data. In each panel is shown the root mean squared error (RMSE) for both types of estimate.

#### Regression analysis

The results in Fig 2 show how poorly fitting a linear model to 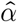 point estimates does, typically overestimating the absolute value of *β* by a large margin. Using MAP estimates of *α* does better here, exhibiting the opposite behaviour of slightly underestimating the absolute value of *β*. However, by cutting out the step of generating point estimates for each pair the GLM retains all pertinent information and accurately recaptures the parameters of the data generating model. It is of course expected that the GLM should be able to discover the correct parameter values, since they were generated by an identical model. What is important is the way the other two models fail by comparison, and of course the fact that we now have the correct likelihood function for such a co-occurrence GLM.

**Figure 2:**
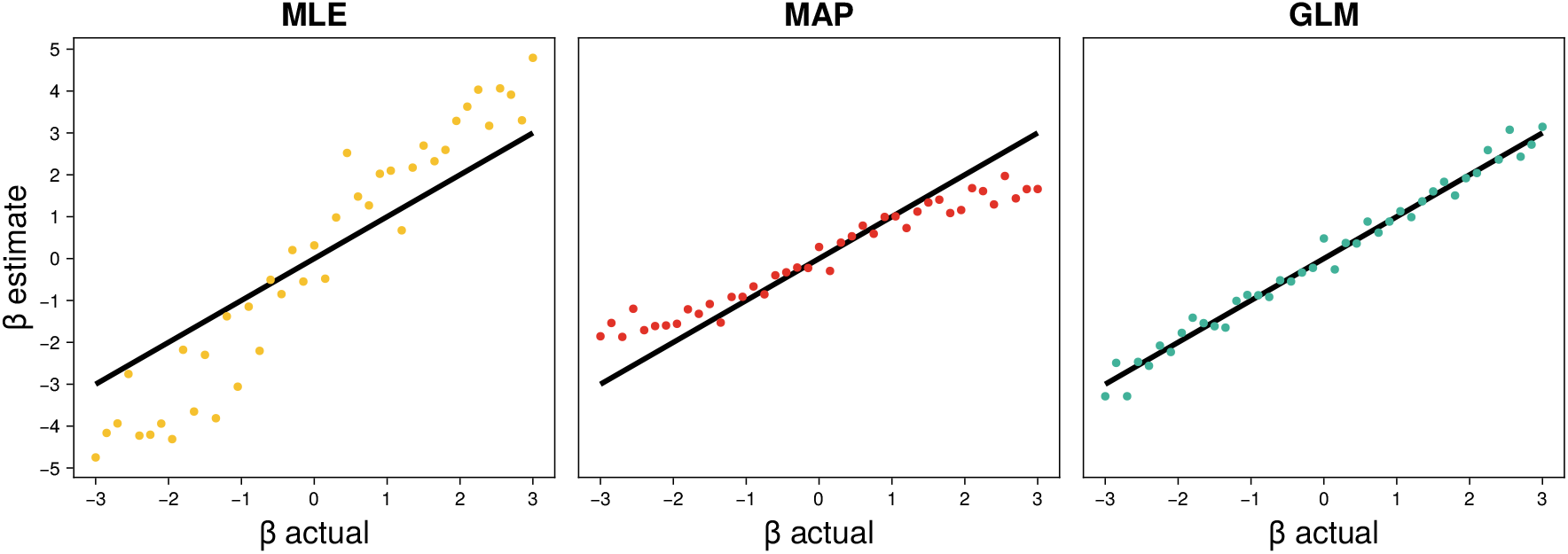
Estimated regression coefficients *β* according to three different methods: fit linear model to 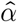 values, fit linear model to MAP estimates of *α*, fit GLM to raw data. Black lines indicate the true *β* values.

### 3.2 Examples with real data

#### Change in co-occurrence affinity owing to trait dissimilarity amongst dung beetles

The first example used occurrence data for dung beetles, along an altitudinal gradient at Serra do Cipó, State of Minas Gerais, Brazil (Nunes et al. 2016). Given the strong abiotic selection imposed by the elevation gradient, I would expect to find greater co-occurrence affinity between more similar beetles. However, there was very little evidence of Gower’s distance (based on both functional guild and mean biomass) having any effect on co-occurrence affinity 0.05 (CI[−0.138, 0.229], PD = 0.703) (Fig 3, top row). By separating trait dissimilarity into functional guild membership and mean biomass, I found that dissimilarity in mean biomass had a significant negative relationship with co-occurrence affinity, with a regression coefficient of −0.342 (CI[−0.621, −0.064], PD = 0.993) (Fig 3, middle row). There was also a significant interaction between the two traits - interaction strength = 0.562 (CI[0.273, 0.879], PD > 0.999), implying that the negative relationship between biomass dissimilarity and co-occurrence affinity is only apparent *within* functional guilds, since the combined effect of biomass and the interaction term (i.e., *β*_biomass_+*β*_biomass_in_different_functional_guild_) was 0.223 (CI[−0.097, 0.556], PD = 0.912). In fact, the effect was so completely confined to within functional guild that the weak evidence of increased affinity with functional guild dissimilarity: *β* = 0.176 (CI[−0.2, 0.541], PD = 0.827) combined with the fact that most species pair were not from the same functional guild led to the slightly positive estimate of the effect of Gower’s distance on affinity. Thus, I present an alternative analysis where I consider only the impact of biomass dissimilarity between species of the same functional guild on co-occurrence affinity (Fig 3, bottom row) - resulting in a slightly increased absolute effect: −0.42 (CI[−0.729, −0.11], PD = 0.997). These results fit the expected pattern of similar species being found together across an environmental gradient but also highlight the possibility of such effects being nuanced in such a way that a combination of expert knowledge and flexible inference method such as the one proposed here may be needed in order to detect them.

**Figure 3:**
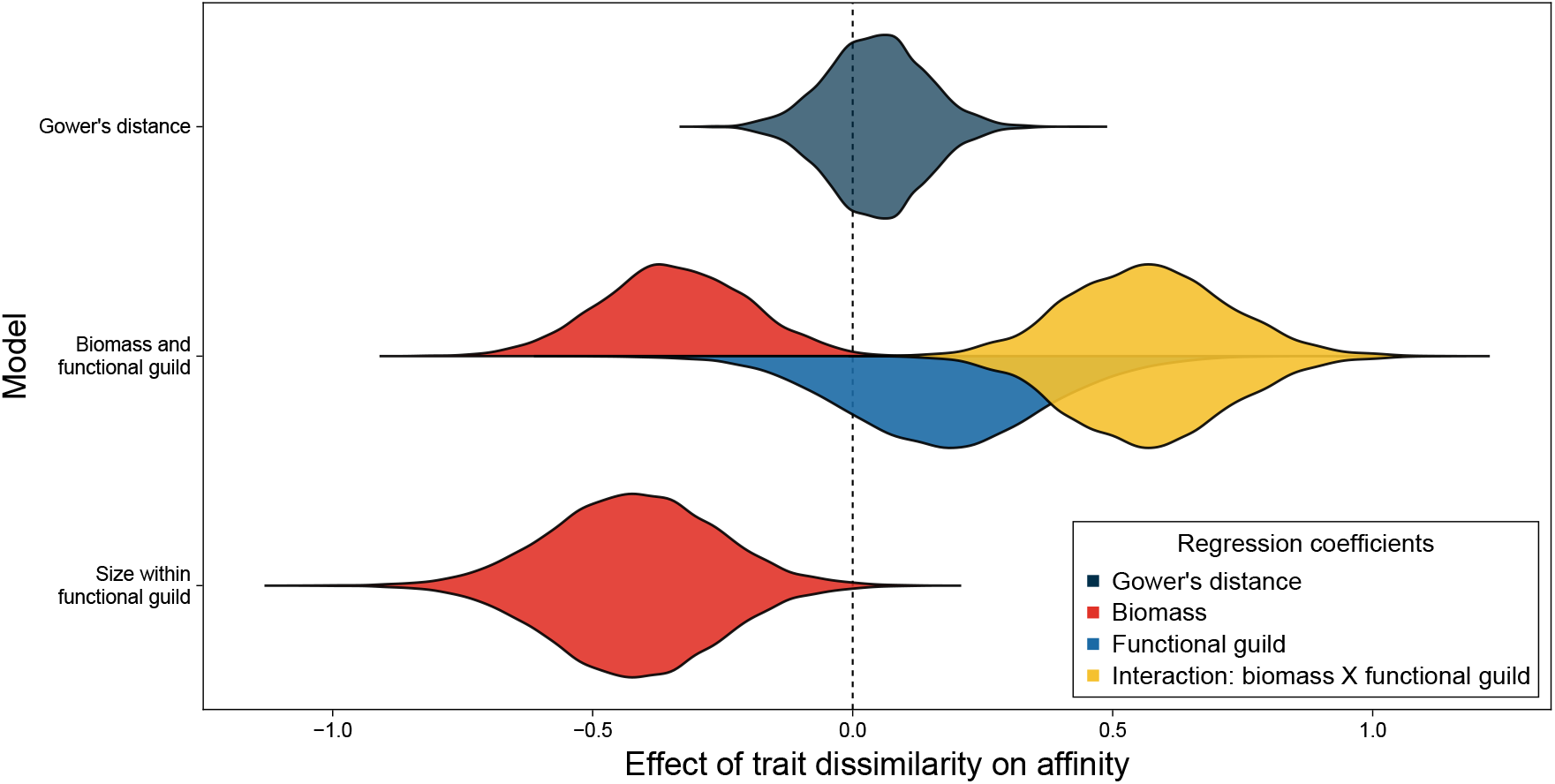
Posterior estimates of change in co-occurrence affinity owing to trait dissimilarity amongst dung beetles, along an altitudinal gradient at Serra do Cipó, State of Minas Gerais, Brazil.

#### Change in co-occurrence affinity owing to trait dissimilarity and relatedness amongst ants

The extent to which functional trait similarity and phylogenetic relatedness lead to taxa co-occurring or being overdispersed are expected to change depending on the geographic scale and (in the case of relatedness) phylogenetic scale under investigation (Webb et al. 2002). Thus, for the next example I made use of ant species occurrence data from both the global scale (Economo et al. 2018) and a smaller scale data set from two nature reserves near Hong Kong (Wong et al. 2021). First, I used mean values for a range of traits (body size, leg length, head width, mandible length, pronotum width and scape length) measured at the Hong Kong sites to see how trait dissimilarity affected affinity at both geographic scales. In general, there was a poor match between the results at different scales. Body size was the only trait that appeared to have the same impact at both scales, with weak evidence of more dissimilar species tending to positively co-occur - *β* = 0.039 (CI[−0.1, 0.176], PD = 0.706) and 0.097 (CI[−0.045, 0.242], PD = 0.908) for local and global scales respectively. The only strong evidence of an effect on affinity in either direction were a negative effect of pronotum width dissimilarity at the global scale: −0.221 (CI[−0.429, −0.013], PD = 0.981), and of head width at the local scale −0.259 (CI[−0.472, −0.042], PD = 0.992) (Fig 4 a).

**Figure 4:**
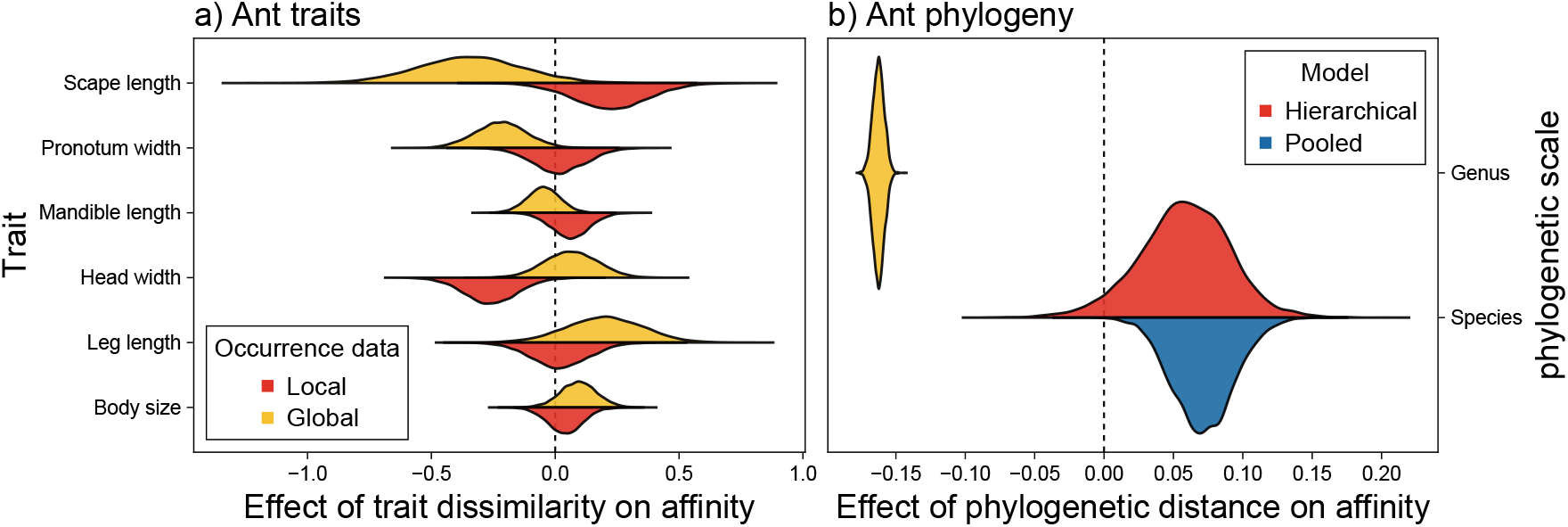
Posterior estimates of change in co-occurrence affinity between ant species owing to a) trait dissimilarity, b) phylogenetic distance.

For the next analysis I used the phylogenetic tree from the global data set (Economo et al. 2018) to ask how phylogenetic distance affects affinity at both species and genus level. The species level analysis investigated the effect of phylogenetic distance on affinity *within* genus (as with the above example of biomass within functional guild). However, with 61 genera to analyse I was also able to employ a hierarchical model with random slopes for each genus. (This modification is simple within the proposed modelling framework and is an option available in the software package `CooccurrenceRegression.jl`). Here, again different scales yielded different results. There was strong evidence for a negative effect of phylogenetic distance on affinity at the genus level: −0.162 (CI[−0.171, −0.155], PD > 0.999) and of a positive relationship at the species level in the non-hierarchical (pooled) model: 0.071 (CI[0.03, 0.115], PD > 0.999). However, there was less certainty in the estimate from the hierarchical model: 0.059 (CI[−0.01, 0.12], PD = 0.955) (Fig 4 b). Here, by treating co-occurrence analysis as just another Bayesian GLM it was straightforward to employ the specific model structure desired to answer the exact question I was interested in - namely a hierarchical structure to measure the effect of relatedness on affinity *within* genus. In this case, while it is reassuring that the pooled model and hierarchical model yield qualitatively similar results, the hierarchical model provides the more reliable results as it naturally accounts for clusters (here genera) in the data (McElreath 2018).

#### Bacterial co-occurrence networks derived from cystic fibrosis patient sputum samples

The final example returns to the issue of simply analysing pairwise co-occurrence relationships, without any explanatory variables. Here, I constructed microbial co-occurrence networks from microbiome data derived from cystic fibrosis patients’ sputum samples (Quinn et al. 2019) using the pairwise affinity estimates (median of posterior MCMC samples) but also retaining and reporting a measure of confidence in those estimates - probability of direction (PD). Fig 5 shows the inferred co-occurrence relationships between the 50 most prevalent genera in the data set.

**Figure 5:**
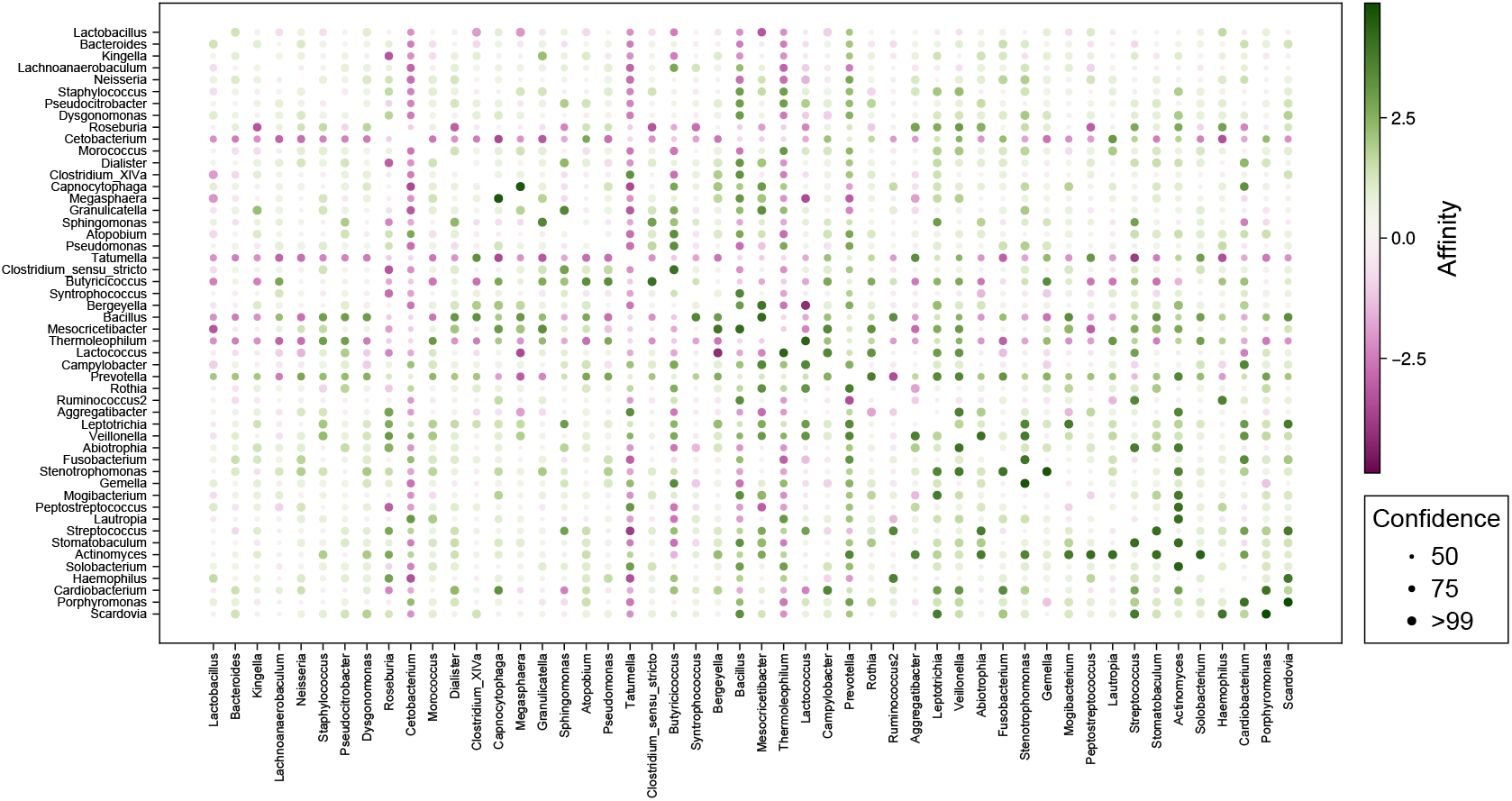
Affinity estimates between the 50 most prevalent genera across a set of cystic fibrosis patient sputum samples. Confidence estimates convey confidence that the affinity between the two genera is at least in the same direction as the point estimate.

Fig 6 Shows an alternative representation - networks constructed from only those genus pairs for which there is over 97.5% confidence in the direction of their affinity. In both representations it can be seen that genus pairs with higher absolute values for co-occurrence affinity do not always have higher confidence in their affinity estimates, though confidence and (absolute) affinity are clearly correlated.

**Figure 6:**
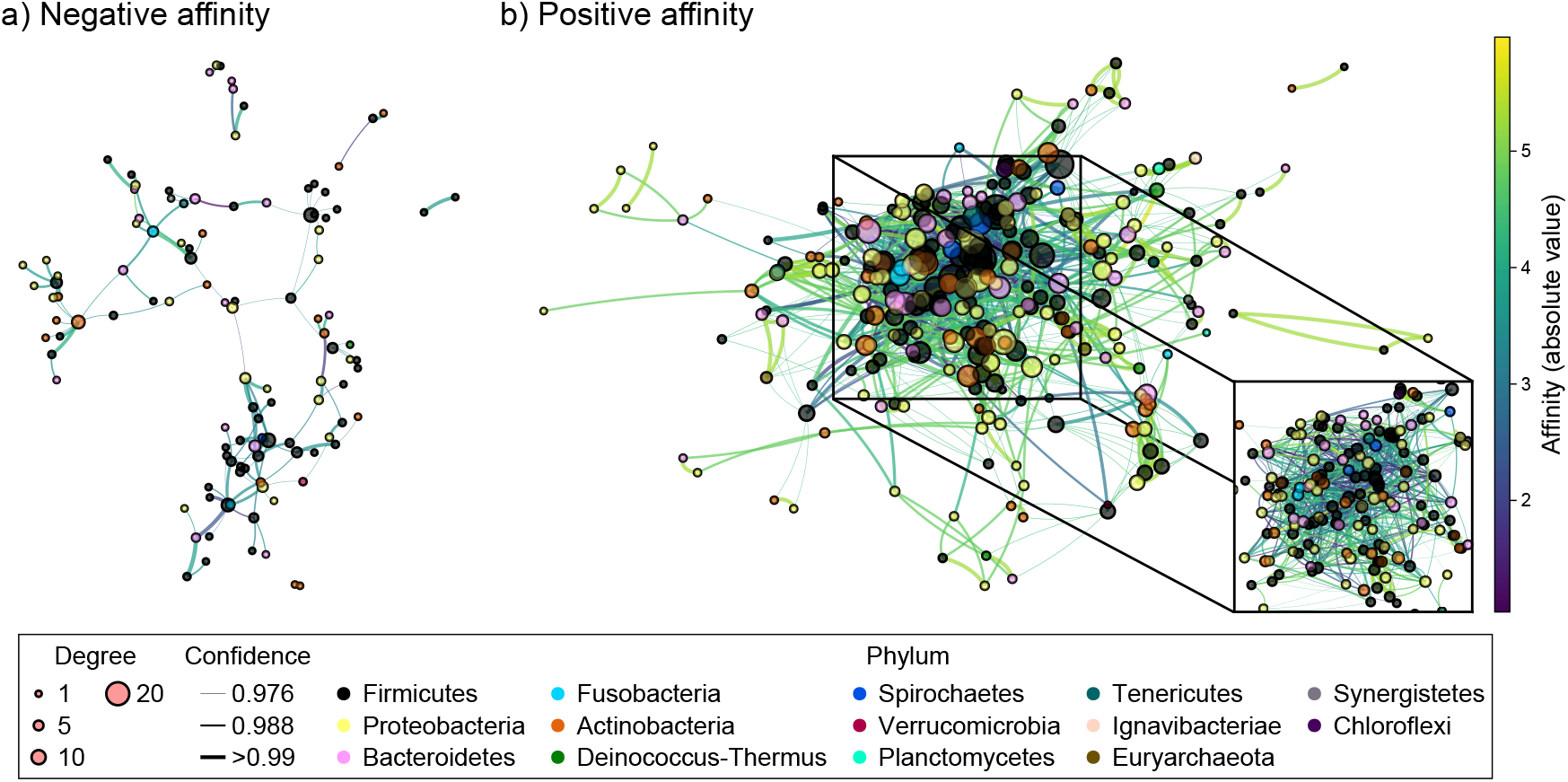
Microbial co-occurrence networks across a set of cystic fibrosis patient sputum samples. a) Negative affinity estimates. b) positive affinity estimates. Each vertex represents a single genus, with its size representing its *degree* (the number of genera it is linked to) and it’s colour representing its phylum. Confidence estimates convey confidence that the affinity between the two genera is at least in the same direction as the point estimate. Only genus pairs with greater than 97.5% confidence in the direction (positive/negative) of their affinity are included. Panel b includes an inset plot of the largest cluster of genera, showing only those links between genera contained in the cluster, with both the edges and vertices of the network reduce in scale for clarity. Here, we can see that these genera are linked together by many relatively small affinity estimates.

### 3.3 CooccurrenceRegression.jl

The main reason behind this work was to communicate a general methodological framework for the benefit of ecologists. In order to simplify the use of this framework I also present the Julia (Bezanson et al. 2017) package CooccurrenceRegression.jl, which is built on top the probabilistic programming language (PPL) Turing.jl (Ge, Xu, and Ghahramani 2018). With this package one can recreate the single explanatory variable (Gower’s distance) model from the dung beetles example for *N* species across *M* sites as follows:

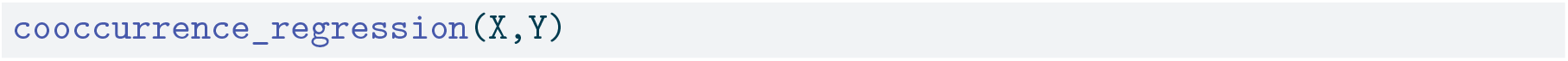

where *X* is an *M* × *M* array of dyadic explanatory variables e.g. a distance matrix and *Y* is an *N* × *M* presence absence matrix with rows of sites and columns of species. Additionally, one could replicate the second model from the dung beetle example by supplying a vector of explanatory matrices:

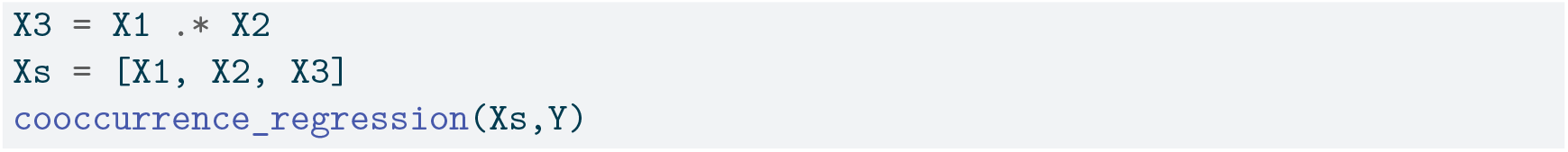

Lastly, hierarchical models can be run as follows

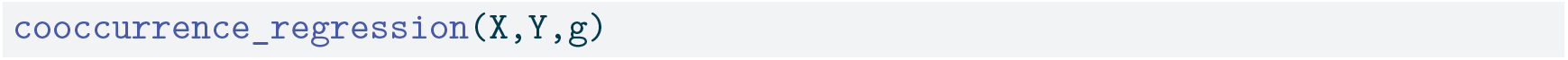

where *g* is a vector of length *M* assigning each species to a particular group.

More detail on changing priors and inference parameters can be found in the package repository https://github.com/EvoArt/CooccurrenceRegression.jl.

### 3.4 Discussion

Here, I have shown how to construct a Bayesian dyadic GLM for the analysis of co-occurrence data. This builds on the work of Mainali et al. (2022) as well as Veech (2013) and Griffith et al. (2016). The identification of Fisher’s noncentral hypergeometric distribution (or mathematically equivalent formulations) as the correct distribution for modelling co-occurrence led first to null model approaches to co-occurrence analysis (Griffith, Veech, and Marsh 2016; Veech 2013), then to a useful co-occurrence metric (Mainali et al. 2022) and now to a general model capable of analysing raw co-occurrence data as a response variable even when data points are not independent (which will generally be the case). It should be noted that the co-occurrence relationships discussed here and in the works cited above are probabilistic in nature, i.e., I do not assume that either a high or low affinity between a specific pair of species implies significant ecological interaction. While co-occurrence analysis does not give the researcher direct access to ecological interaction data, it does have some baring (Cazelles 2024) and may be used as an initial screening to identify likely interactions.

Part of the motivation for this work was the failure to recapture known regression coefficients when fitting linear models to pairwise affinity estimates (Fig 2). There may be other ways of combating this failure. For example, the removal of data points for which we have low confidence may be an option. For instance, if *N* = 30, *mA* = 29 and *mB* = 1, then a *k* of 1 tells us very little, since our null expectation is that *k* will very likely = 1. This will lead to a high affinity estimate but with a wide confidence interval and a high p-value. Using a cut-off threshold e.g. only using data for which p < 0.05 may lead to better results. However, these data points are now *missing* from further analyses. The potential pitfalls involved in dealing with missing data are numerous (Kang 2013), but it is unnecessary to risk the possible bias associated with systematically removing data points when the Bayesian analysis framework naturally accounts for differing levels of confidence between data (McElreath 2018).

The method proposed here is as flexible as any Bayesian GLM and can thus be adjusted to fit the specific questions and modelling assumptions of the researcher. Here, I have shown the applicability of this approach to analysing the relationship between trait dissimilarity or phylogenetic distance and co-occurrence, as well as constructing co-occurrence networks. In doing so, I have needed to use hierarchical models and interacting explanatory variables to get accurate results. Many more model structures can be employed, as well as many more applications inside and outside of ecology e.g. social networks in the humanities and gene co-occurrence relationships in genetics. For simple models with a single presence/absence matrix as response and one or more matrices of explanatory variables I have developed the Julia (Bezanson et al. 2017) package CooccurrenceRegression.jl. However, it will serve many researchers to consult accessible texts on probablistic programming (McElreath 2018), read the source code of the package, and develop their own models to suit their specific research question.

While the focus here was on regression, retaining information on both confidence/uncertainty and estimated effect size will also be important for downstream analyses of co-occurrence neworks e.g. generating random networks and analysing network metrics for each. However, with the approach used here, the minimum confidence in any interaction is 50% (unless an effect size threshold is used e.g. the probability of an absolute affinity value >1). I used probability of direction as a measure of confidence in the visualisations, because researchers are often interested in the binary classification of positive vs negative. However, other measures (e.g. 1/(width of 95% credible interval)) may also be used. Future work should investigate the use of sparsity inducing spike and slab priors and their approximations (Castillo, Schmidt-Hieber, and Van der Vaart 2015), to model the assumption that many species will not interact in any meaningful way. Although, assumption may or may not be valid for the affinity metric, baring in mind that affinity measures a statistical likelihood for species to co-occur, and does not directly measure biotic interactions.

The method proposed here provides ecologist with an important new tool for the analysis of co-occurrence, and in particular discovering the relationships between co-occurrence and other variables e.g. phylogenetic distance, which is an active area of research (Goberna et al. 2019) and has been the subject of much research effort over the past few decades (Webb et al. 2002) but now has an analysis framework based on the simple application of probability theory (McElreath 2018) with correct modelling of co-occurrence probabilities (Griffith, Veech, and Marsh 2016; Veech 2013; Mainali et al. 2022) while accounting for non-independence of data in matrices of pairwise species measurements (Clarke, Rothery, and Raybould 2002; Gompert et al. 2014).

## Code and data availabilty

All code and associated with the manuscript can be fount at https://github.com/EvoArt/bayesian-affinity.

